# Synthetic oxepanoprolinamide iboxamycin is highly active against human pathogen *Listeria monocytogenes*

**DOI:** 10.1101/2022.02.28.482263

**Authors:** Tetiana Brodiazhenko, Kathryn Jane Turnbull, Kelvin J.Y. Wu, Hiraku Takada, Ben I.C. Tresco, Tanel Tenson, Andrew G. Myers, Vasili Hauryliuk

## Abstract

Listeriosis is a dangerous food-borne bacterial disease caused by the Gram-positive Bacillota (Firmicute) bacterium *Listeria monocytogenes.* In this report, we show that the synthetic lincosamide iboxamycin is highly active against *L. monocytogenes* and can overcome the intrinsic lincosamide resistance mediated by VgaL/Lmo0919, a member of ABCF ATPase resistance determinants that act by directly removing the antibiotic from the ribosome. While iboxamycin is not bactericidal against *L. monocytogenes,* it displays a pronounced postantibiotic effect, which is a valuable pharmacokinetic feature. Experiments in *L. monocytogenes* infection models are necessary to further assess the potential of iboxamycin as a novel drug for treatment of listeriosis. We demonstrate that VmlR ARE ABCF of Bacillota bacterium *Bacillus subtilis* grants significant (33-fold increase in MIC) protection from iboxamycin, while LsaA ABCF of *Enterococcus faecalis* grants an 8-fold protective effect. Furthermore, the VmlR-mediated iboxamycin resistance is cooperative with that mediated by the Cfr 23S rRNA methyltransferase resistance determinant, resulting in up to a 512-fold increase in MIC. Therefore, emergence and spread of ABCF ARE variants capable of defeating next-generation lincosamides in the clinic is possible and should be closely monitored.

## Introduction

Lincosamides constitute an important class of antibiotics used both in veterinary and human medicine [1]. These compounds inhibit protein synthesis by binding to and compromising the enzymatic activity of the peptidyl transferase centre (PTC) of the ribosome [2–5], resulting in bacteriostasis [6]. Representatives of this antibiotic class share a common architecture and are typically comprised of a 4’-substituted L-proline residue connected via an amide bond to a unique S-glycosidic aminosugar moiety (**Figure 1A,B**). The first lincosamide to be discovered, lincomycin (**Figure 1A**), is a natural product produced by *Streptomyces lincolnensis* ssp. *lincolnensis* and is active against streptococcal, pneumococcal and staphylococcal infections [7]. Its semi-synthetic derivative, clindamycin (**Figure 1B**), can be produced via a one-step stereoinvertive deoxychlorination of lincomycin [8]. Clindamycin is more potent than lincomycin and is currently the lincosamide of choice for human medicine [9]. Like lincomycin, clindamycin is mostly active against Gram-positive but not Gram-negative bacteria, which restricts the spectrum of its applications [10]. A *cis*-4-ethyl-L-pipecolic acid amide of clindamycin, pirlimycin, has a similar spectrum of antibacterial activity [11, 12] and is approved for veterinary applications in the United States and European Union. Finally, a recently developed semisynthetic derivative of lincomycin (‘compound A’) was shown to be able to overcome clindamycin resistance in *Staphylococcus aureus* mediated by ribosomal RNA (rRNA) methylation by ErmA and ErmB antibiotic resistance determinants [13].

**Figure 1.**
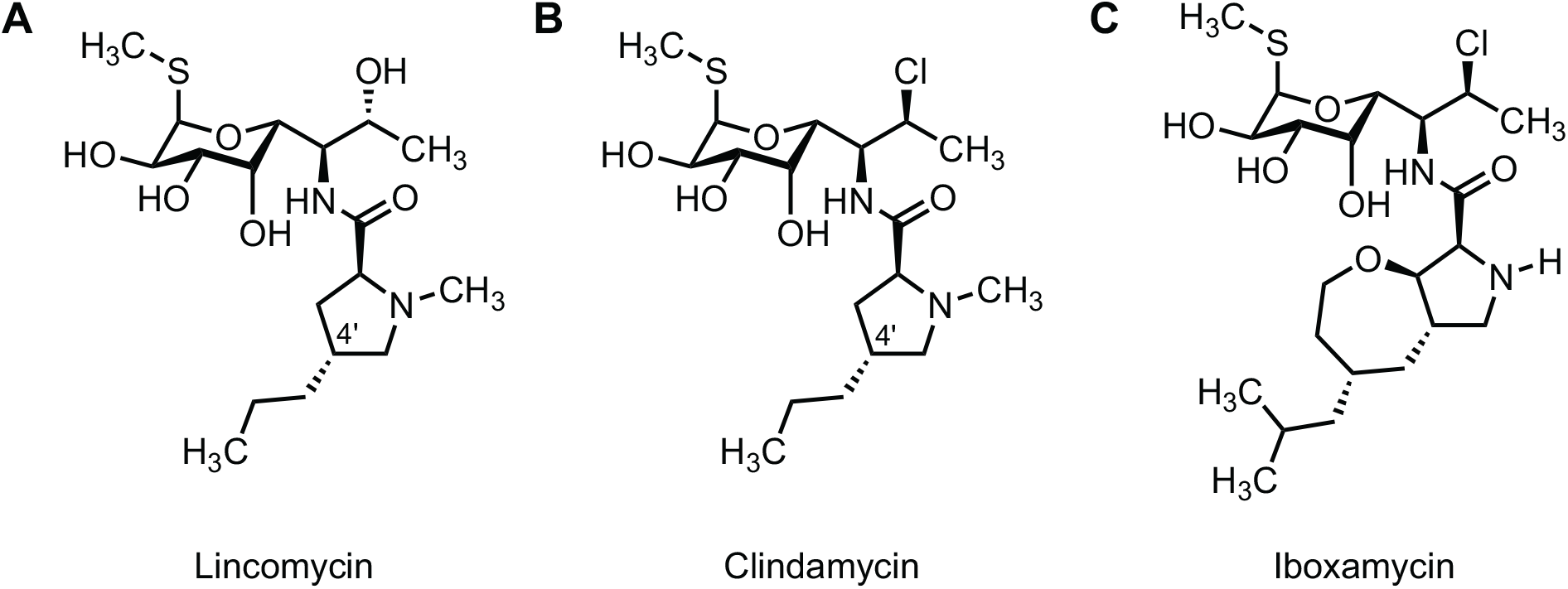
Chemical structures of lincosamide antibiotics lincomycin (A), clindamycin (B) and iboxamycin (C).

Iboxamycin (**Figure 1C**) is a newly developed lincosamide with an exceptionally broad spectrum of antibacterial activity [14]. Featuring a fully synthetic, bicyclic oxepanoprolinamide aminoacyl fragment, iboxamycin improves upon previous lincosamides in its activity against both Gram-positive and Gram-negative pathogens [14]. Iboxamycin was found to be more potent than clindamycin against Gram-positive pathogens and overcomes lincosamide resistance mediated by rRNA modification by Erm and Cfr 23S rRNA methyltransferases, both of which are highly clinically important and widespread antibiotic resistance determinants [15–18]. While Cfr grants strong protection against clindamycin in clinical isolates of *Staphylococcus aureus* and *Staphylococcus epidermidis* (MIC >128 μg/mL), it confers only moderate resistance against iboxamycin (MIC of 2-8 μg/mL compared to 0.06 μg/mL for *cfr-*strains) [14]. Importantly, iboxamycin is also highly active against *Enterococcus faecalis* (MIC 0.06 μg/mL as compared to 16 μg/mL for clindamycin) – a species that is intrinsically resistant to ‘classical’ lincosamides as it encodes the LsaA antibiotic resistance (ARE) factor in its chromosomal genome [19], a member of the ABCF ATPase protein family that includes multiple resistance factors [20–22]. LsaA provides resistance against pleuromutilin, lincosamide and streptogramin A (PLS_A_) antibiotics by displacing the drug from the ribosome [23], acting similarly to other ARE ABCFs [24–27]. As evident from the 96- to 256-fold higher sensitivity to clindamycin and lincomycin in a Δ*lsaA E. faecalis* strain as compared to *E. faecalis* ectopically expressing LsaA [23], LsaA is a potent lincosamide resistance determinant. The high sensitivity of *E. faecalis* to iboxamycin suggests that this compound has the potential to overcome resistance mediated by other ARE ABCFs as well.

Listeriosis is a dangerous food-borne bacterial disease caused by the Gram-positive Bacillota (formerly: Firmicute) bacterium *Listeria monocytogenes,* which infects people through contaminated meat, fish and dairy products [28, 29]. While it is a relatively rare infection that mainly affects people with weakened immune systems, or who are pregnant [30], the majority of listeriosis cases require hospitalisation and mortality rates can be as high as 20-30% even with antibiotic treatment [31, 32]. Antibiotic treatment options for *L. monocytogenes* infections include cell wall synthesis disruptors ampicillin and vancomycin, folic acid synthesis inhibitors sulfamethoxazole and trimethoprim, and protein synthesis inhibitors, such as gentamicin and azithromycin [33]. *L. monocytogenes* strains reported in recent years are often resistant to clindamycin, with the resistant fraction ranging from 29% to 76%, depending on the collection [34–37], thus excluding clindamycin as a viable option for treatment of *L. monocytogenes* infections. Importantly, just as *E. faecalis* encodes the ABCF ATPase LsaA, *L. monocytogenes* encodes the ARE ABCF PLS_A_ resistance factor VgaL/Lmo0919 in its core genome [38]. As with LsaA, VgaL operates on the ribosome [23], and loss of VgaL results in increased sensitivity to lincosamides, with the Δ*lmo0919 L. monocytogenes* strain being 8- to 16-fold more sensitive to lincomycin as compared to the isogenic wild type [23]. Finally, a model Bacillota, *B. subtilis,* also encodes an ARE ABCF PLS_A_ resistance factor – VmlR [27, 39].

In this report, using lincomycin and clindamycin as reference compounds, we i) characterised the efficacy of iboxamycin against *L. monocytogenes,* ii) probed its ability to specifically counter resistance mediated by ABCFs *L. monocytogenes* Lmo0919, *E. faecalis* LsaA and *B. subtilis* VmlR, iii) characterised its bactericidal/bacteriostatic mechanism of action and, finally, iv) assessed the strength of its post-antibiotic effect (PAE).

## Results

### L. monocytogenes *is highly sensitive to iboxamycin despite VgaL/Lmo0919 ABCF resistance factor*

To test the lincosamide sensitivity of *L. monocytogenes* we used two widely-used wild-type strains, both belonging to serovar 1/2a: EGD-e [40] and 10403S, a streptomycin-resistant variant of 10403 [41]. The two wild types are genomically distinct, e.g. the virulence master-regulator PrfA is overexpressed in EGD-e and the prophage content differs between the two strains [42]. In additional to the two wild types, we also tested a *L. monocytogenes* EDG-e derivative that was genomically modified to abrogate the expression of VgaL/Lmo0919 PLS_A_ resistance factor (EDG-e Δ*lmo0919*) [23].

Both wild-type *L. monocytogenes* strains are dramatically more sensitive to iboxamycin (MIC of 0.125-0.5 μg/mL) as compared to clindamycin (MIC of 1 μg/mL) and lincomycin (MIC of 2-8 μg/mL) (**Table 1**). In agreement with the higher sensitivity of Δ*lmo0919* EDG-e to lincomycin [23], this strain is 2-8-fold more sensitive to iboxamycin than the corresponding wild type. This indicates that while VgaL does confer some protection from iboxamycin, the high potency of the synthetic antibiotic would likely allow the drug to overcome resistance in clinical settings. A likely explanation is that increased affinity of the synthetic drug for the ribosome renders antibiotic displacement by ABCF ATPases inefficient.

**Table 1.** Broth microdilution Minimum inhibitory concentration (MIC) testing of lincosamide antibiotics against *L. monocytogenes, E. faecalis* and *B. subtilis* strains. In the case of *L. monocytogenes* strains, MIC testing was carried out in MH-F broth and growth inhibition was scored after 48 hours incubation at 37 °C. *E. faecalis* MIC testing was carried out in BHI broth supplemented with 2 mg/mL kanamycin (to prevent *lsa* revertants), 0.1 mg/mL spectinomycin (to maintain the pCIE_spec_ plasmid), 100 ng/mL of cCF10 peptide (to induce expression of LsaA protein). *B. subtilis* MIC testing was carried out in either LB medium or LB supplemented with 1 mM IPTG to induce expression of either VmlR or Cfr protein, and growth inhibition was scored after 16-20 hours at 37 °C.

| Species / Strain | Antibiotic MIC, µg/mL |  |  |
| --- | --- | --- | --- |
|  | Lincomycin | Cindamycin | Iboxamycin |
| *L. monocytogenes* 10403S | 4-8 | 2 | 0.125-0.25 |
| *L. monocytogenes* EDG-e | 8 | 1-2 | 0.125-0.5 |
| *L. monocytogenes* EDG-e Δlmo0919 | 0.25-1 | 0.125-0.5 | 0.0625 |
| *E. faecalis* ΔlsaA pCIE<sub>spec</sub> | 0.125 | 0.15 | 0.0625 |
| *E. faecalis* ΔlsaA pCIE<sub>spec</sub> LsaA | 16-32 | 16 | 0.5 |
| *B. subtilis* wt 168 | 80 | 4 | 2 |
| *B. subtilis* ΔvmlR | 2.5 | 0.125 | 0.06 |
| *B. subtilis* ΔvmlR thrC::P<sub>hy-spank</sub>-vmlR (IPTG: 1 mM) | 160 | 8 | 4 |
| *B. subtilis* thrC::P<sub>hy-spank</sub>-cfr (IPTG: 1 mM) | >640 | 640 | 16-32 |
| *B. subtilis* ΔvmlR thrC::P<sub>hy-spank</sub>-cfr (IPTG: 1 mM) | >640 | 320 | 2 |

Importantly, expression of Lmo0919 is not constitutive: it is elicited by antibiotic-induced ribosomal stalling on the regulatory short open reading frame upstream of the *lmo0919* gene [38]. Therefore, the difference in iboxamycin sensitivity between wild-type and Δ*lmo0919* EDG-e strains reflects both the ability of Lmo0919 to protect the ribosome from the antibiotic as well as the efficiency of iboxamycin-mediated induction of Lmo0919. To deconvolute these two effects, we used engineered strains that allow for ectopic inducible expression of ABCF in the following experiments.

### E. faecalis *ABCF LsaA grants a moderate protective effect against iboxamycin*

To test the ability of other ABCF PLS_A_ resistance factors to confer resistance to iboxamycin, we compared a pair of *E. faecalis* strains: one lacking the chromosomally-encoded LsaA (*DlsaA* pCIEspec) and the other allowing cCF10-peptide-inducible expression of LsaA (*DlsaA* pCIEspec LsaA) [23]. Using this experimental set up, we could specifically assess the ability of LsaA to protect the strain from lincosamides. While expression of LsaA dramatically increases resistance to clindamycin and lincomycin (96- to 256-fold, respectively), it results in a mere 8-fold protective effect against iboxamycin (MIC of 0.0625 and 0.5 μg/mL, respectively) (**Table 1**), demonstrating that iboxamycin can also largely overcome LsaA-mediated resistance.

### B. subtilis *ABCF VmlR acts cooperatively with rRNA methyltransferase Cfr to grant significant protection against iboxamycin*

Next we tested a set of *B. subtilis* strains: wild-type 168 *B. subtilis, DvmlR* (VHB5) as well as a *DvmlR* strain in which VmlR is expressed under the control of IPTG-inducible *Phy-spank* promotor (VHB44) [43] (**Table 1**). Disruption of *vmlR* results in a 33-fold increase in iboxamycin sensitivity (MIC of 2 and 0.06 μg/mL, respectively), and resistance is restored upon ectopic expression of VmlR (MIC of 4 μg/mL, 2-fold increase over the wild-type levels). The iboxamycin sensitivity of Δ*lmo0919 L. monocytogenes* EDG-e and *DvmlR B. subtilis* is near-identical, indicating that the 16-/4-fold difference in iboxamycin sensitivity between wild-type *L. monocytogenes* and *B. subtilis* is due to the different efficiency of resistance granted by Lmo0919 and VmlR respectively.

Importantly, VmlR loss results in the same *relative* increase in sensitivity to all lincosamides tested – iboxamycin, clindamycin and lincomycin; 32-33-fold – regardless of the potency of the lincosamide (**Table 1**). This suggests that if the affinity of iboxamycin to the target were to be decreased by, for instance, rRNA modification, direct target protection by the ABCF could cooperatively lead to high levels of resistance. To probe this hypothesis, we have characterised the lincosamide sensitivity of *B. subtilis* strains that express Cfr 23S rRNA methyltransferase under the control of IPTG-inducible *Phy-spank* promotor, either in the presence or absence of the chromosomally-encoded VmlR. Ectopic expression of Cfr in *vmlR+ B. subtilis* effected a cooperative resistance to iboxamycin, resulting in MICs of 16-32 μg/mL as opposed to 2 μg/mL when either of these resistance determinants are expressed individually (**Table 1**). As expected, Cfr also granted high levels of lincomycin and clindamycin resistance when ectopically expressed in both wild-type and *DvmlR* strains (MIC ranging from 320 to excess of 640 μg/mL).

### *Iboxamycin is bacteriostatic against* L. monocytogenes *and displays a strong postantibiotic effect*

Macrolide antibiotics that tightly bind the ribosome and dissociate slowly are bactericidal, while macrolides that dissociate rapidly are bacteriostatic [44]. As with lincomycin and clindamycin, iboxamycin was shown to be bacteriostatic against a panel of bacterial species [14]. However, since effects on *L. monocytogenes* were not assessed in the original report – and the species is highly sensitive to iboxamycin – we tested for potential bactericidal effects of iboxamycin against this pathogen. The three *L. monocytogenes* strains that we used for the MIC measurements – wild-type 10403 and EGD-e as well as ABCF-deficient EDG-e Δ*lmo0919* – were treated with 4x MIC concentration of either iboxamycin, clindamycin and lincomycin for increasing periods of time (from 2 to 24 hours), washed, and then plated on BHI agar plates that contain no antibiotic. The bacterial growth expressed in Colony Forming Units, CFU, was scored after either 24- or 48-hour incubation of plates at 37 °C. When the colony counting was performed after 24 h, we observed potentially bactericidal behaviour of iboxamycin, with almost a two log_10_ drop in CFU after the 10-hour treatment with the antibiotic (**Figure 2A-C**). Importantly, no similar CFU decrease was observed for either clindamycin or lincomycin (**Figure 2A-C**). However, this apparent CFU drop effect of iboxamycin disappeared after 48 h of incubation (**Figure 2D-F**), suggesting slow regrowth rather than cidality and indicative of the so-called postantibiotic effect (PAE) [45, 46].

**Figure 2.**
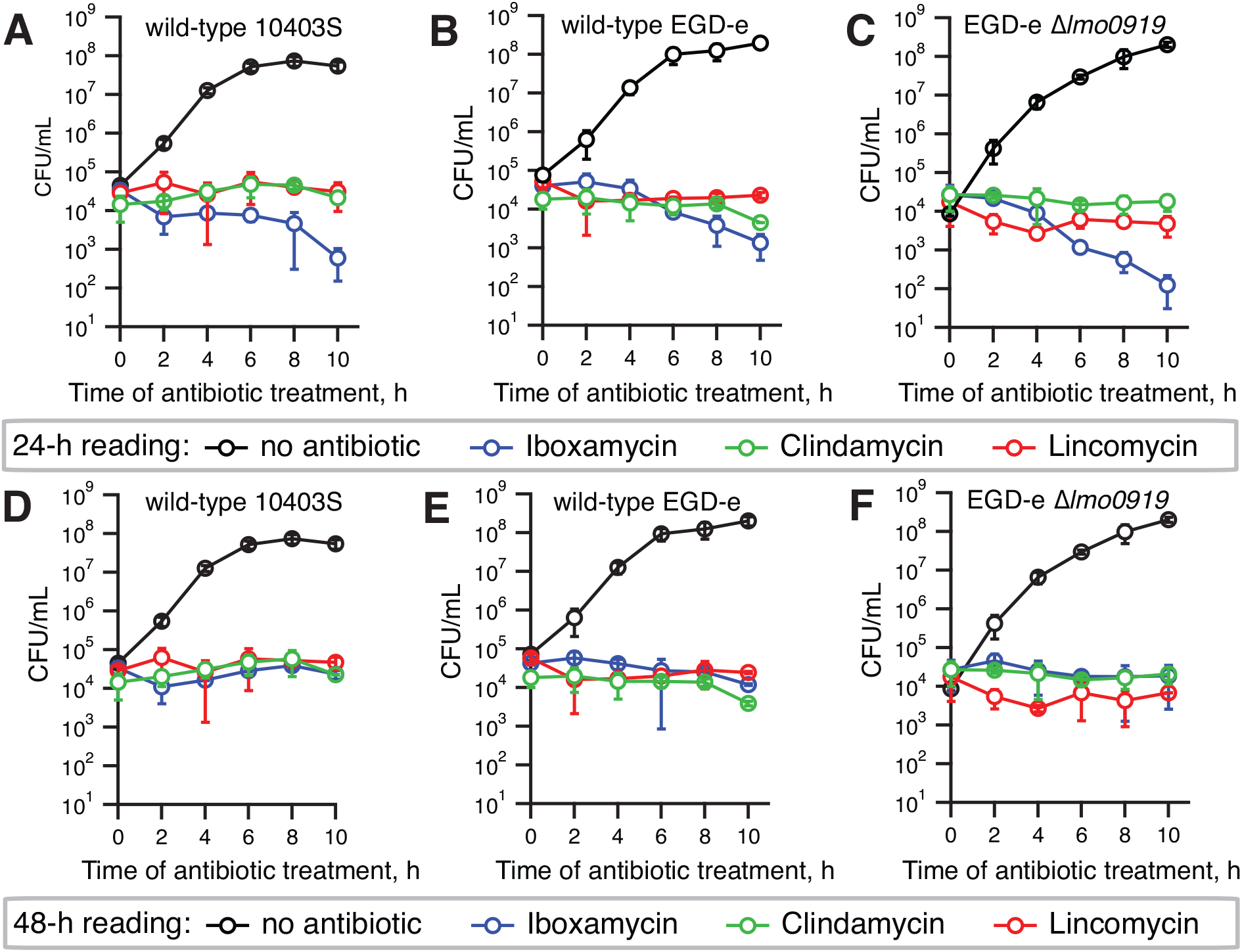
Iboxamycin is bacteriostatic against *L. monocytogenes*. Exponentially growing *L. monocytogenes* type strains; 10403S (**A,D**), EDG-e (**B,E**) or VgaA-deficient EDG-e Δ*lmo0919* (**C,E**) were treated with 4x MIC of either iboxamycin, clindamycin, lincomycin or no antibiotic as control. Cells were harvested at given time points and washed before plating. After 24 (**AC**) or 48 hours (**D-F**) of incubation, colonies were counted to determine CFU/ml. All experiments were carried out in MH-F broth at 37 °C with shaking at 180 rpm, data points are from three biological replicates and standard deviation is indicated with error bars.

PAE is characterised by the time after antibiotic removal where no growth of the treated bacteria is observed. This prolonged action of iboxamycin has been previously noted for *S. aureus* and *E. faecium* [14]. Therefore, we next performed post-antibiotic effect experiments in *L. monocytogenes,* demonstrating that, indeed, iboxamycin displays pronounced PAE, suppressing the growth of the wildtype 10403S and wild-type EGD-e for 6 and 8 hours, respectively (**Figure 3B,C**). Clindamycin demonstrates a weaker PAE against EGD-e (2 hours) and similar PAE against 10403S. No clear PAE is detectible for lincomycin. Compared with the isogenic wild-type, EDG-e Δ*lmo0919* displays similar PAE in the case of clindamycin, and, possibly, somewhat more pronounced PAE in the case of iboxamycin.

**Figure 3.**
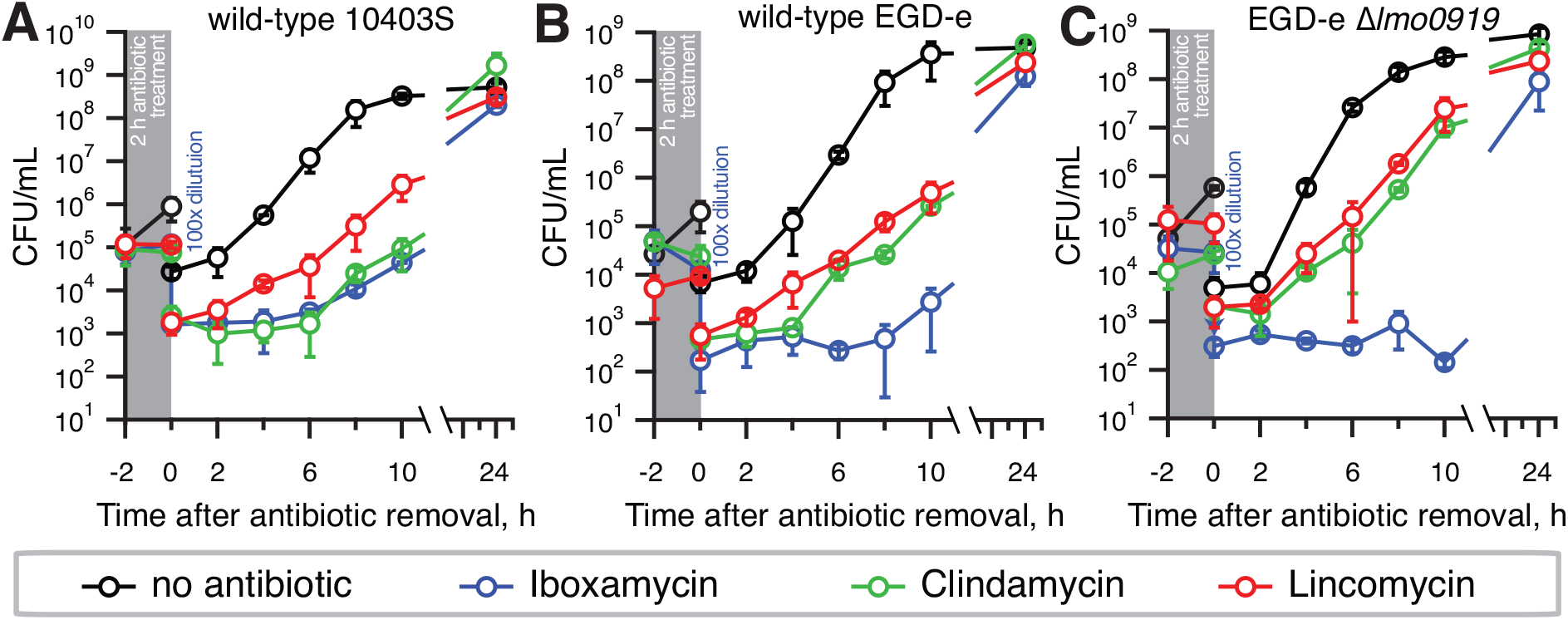
Iboxamycin displays strong postantibiotic effect against *L. monocytogenes.* To determine the time taken for antibiotic treated *L. monocytogenes* strains to resume growth after a two-hour antibiotic treatment, exponentially growing type strains; 10403S (**A**), EDG-e (**B**) or VgaA-deficient EDG-e Δ*lmo0919* (**C**) were treated with 4x MIC of either iboxamycin, clindamycin, lincomycin, or no antibiotic as control, for two hours. Cells were then diluted by 100-fold to remove the antibiotic, and samples taken every two hours subsequently for viability counting. All experiments were carried out in MH-F broth at 37 °C with shaking at 180 rpm, data points are from three biological replicates and standard deviation is indicated with error bars.

## Discussion

In this report we have evaluated the efficiency of the oxepanoprolinamide iboxamycin against *L. monocytogenes.* The antibiotic can largely overcome the intrinsic PLS_A_ resistance of this species that is mediated by the ribosome-associated ATPase VgaL/Lmo0919, and can similarly counteract the intrinsic resistance mediated by ARE ABCF LsaA in *E. faecalis.* ARE ABCF PLS_A_ resistance factors are broadly distributed among bacterial pathogens [20, 22, 47, 48], and therefore the ability of iboxamycin to largely counteract the ABCF-mediated resistance is a valuable feature of the new antibiotic. However, given that *B. subtilis* VmlR does confer significant levels of iboxamycin resistance (33-fold increase in MIC) and is cooperative with the Cfr rRNA methyltransferase resistance determinant, emergence and spread of ABCF ARE variants capable of defeating next-generation lincosamides in the clinic is possible and should be closely monitored.

Furthermore, we demonstrate that iboxamycin displays a strong PAE against *L. monocytogenes,* compromising bacterial re-growth for many hours post antibiotic removal. The PAE is considerably stronger than that of clindamycin while lincomycin displays no PAE. It is possible that the strength of the PAE reflects how tightly the antibiotic binds to the target, the ribosome – and how slowly it dissociates from it. The pronounced PAE suggests that development of even more tight-binding lincosamides could produce effectively bactericidal drugs in the context of infection. Further biochemical studies are necessary to substantiate this hypothesis. Experiments in *L. monocytogenes* infection models are necessary to further assess the potential of iboxamycin as a novel drug for the treatment of listeriosis.

## Materials and methods

### Synthesis of Iboxamycin

Iboxamycin was prepared according to the method reported by Mason *et al.* [49].

### Strains and media

Wild-type *L. monocytogenes* 10403S was provided by Daniel A. Portnoy, wild-type *L. monocytogenes* EGD-e was provided by Jörgen Johansson, construction of *L. monocytogenes* EDG-e Δ*lmo0919* was described earlier [23], *E. faecalis DlsaA* (*lsa*::Kan) strain TX5332 [19] was provided by Barbara E. Murray, *E. faecalis DlsaA* pCIE_spec_ and *E. faecalis DlsaA* pCIE_spec_ LsaA were described earlier [23]. Wildtype 168 *trpC B. subtilis* (laboratory stock) was used. *B. subtilis* strains *trpC DvmlR* (VHB5) and *DvmlR thrC::P_hyspank_-vmlR* (VHB44) were described earlier [27]. To construct *B. subtilis thrC:: P_hyspank_-cfr* (VHB138) and *DvmlR thrC:: P_hyspank_-cfr* (VHB139), a PCR product encoding *Staphylococcus sciuri cfr* gene optimized to *E. coli* codon usage [50] was PCR-amplified from the pBRCfr plasmid using primers VHT25 (5’-CGGATAACAATTAAGCTTAGTCGACTTAAGGAGGTGTGTCTCATGAACTTTAACAACAAAACCAAATAC-3’) and VHT26 (5’-GTTTCCACCGAATTAGCTTGCATGCTCACTGGGAGTTCTGATAGTTACCATACA-3’). The second PCR fragment encoding a kanamycin-resistance marker, a polylinker downstream of the *P_hyspank_* promoter and the *lac* repressor ORF – all inserted in the middle of the *thrC* gene – was PCR-amplified from pHT009 plasmid using primers pHT002_F (5’-GTCGACTAAGCTTAATTGTTATCCGCTCACAATTACACACATTATGCC-3’) and pHT002_R (5’-GCATGCAAGCTAATTCGGTGGAAACGAGGTCATC-3’). The two fragments were ligated using the NEBuilder HiFi DNA Assembly master mix (New England BioLabs, Ipswich, MA) yielding the pHT009-*cfr* plasmid (VHp439) which was used to transform either wild-type 168 *trpC2* or *DvmlR* (VHB5) strain. Selection for kanamycin resistance yielded the desired VHB138 and VHB139 strain.

Growth assays, MIC, cidality and post antibiotic effect assays with *L. monocytogenes,* were performed in MH-F broth, *E. faecalis* MIC assays were performed in BHI broth and *B. subtilis* MIC assays were performed in LB broth. The media was prepared as per European Committee on Antimicrobial Susceptibility Testing (EUCAST) guidelines (https://www.eucast.org/fileadmin/src/media/PDFs/EUCAST_files/Disk_test_documents/2020_manuals/Media_preparation_v_6.0_EUCAST_AST.pdf) and contained 95% Mueller-Hinton broth (MHB) media (Sigma, Lot# BCCB5572), 5% lysed horse blood (defibrinated 50% stock, Hatunalab AB Cat. N° 139) and 20 mg/mL β-NAD (Sigma, Lot# SLCD5502). Prior to use the 50% horse blood stock was freeze thawed five times and clarified via centrifugation twice for 30 minutes at 18,000 rpm at 4 °C and then filtrated using 0.2 μm membrane filter, aliquoted and stored at −20°C. Solid agar plates were prepared from BHI broth media (VMR, Lot# G0113W) supplemented with 1% (final concentration) agar.

### Liquid growth assays

*L. monocytogenes* was pre-grown on BHI agar plates at 37 °C for 48 hours. Individual fresh colonies were used to inoculate 2 mL of MH-F broth in 15 mL round bottom tubes, which were then incubated overnight at 37 °C with shaking at 180 rpm. The overnight cultures were diluted then with MH-F broth to final OD_600_ of 0.005 and incubated for 8 hours in a water bath shaker (Eppendorf™ Inova™ 3100 High-Temperature) at 37 °C with shaking at 160 rpm. bacterial growth was monitored by OD_600_ measurements every 30 minutes.

### Antibiotic susceptibility testing

The Minimum Inhibitory Concentration (MIC) antibiotic sensitivity testing was performed according to EUCAST guidelines (http://www.eucast.org/ast_of_bacteria/mic_determination), as described earlier [23].

*L. monocytogenes* strains were grown in MH-F broth inoculated with 5 x 10^5^ CFU/mL (OD_600_ of approximately 0.0015) with increasing concentrations of antibiotics. After 24-48 hours of incubation at 37 °C without shaking, the presence or absence of bacterial growth was scored by eye.

*E. faecalis* strains were grown in BHI media supplemented with 2 mg/mL kanamycin (to prevent *lsa* revertants), 0.1 mg/mL spectinomycin (to maintain the pCIE_spec_ plasmid), 100 ng/mL of cCF10 peptide (to induce expression of LsaA protein) as well as increasing concentrations of antibiotics, was inoculated with 5 × 10^5^ CFU/mL (OD_600_ of approximately 0.0005) of *E. faecalis DlsaA (lsa::Kan)* strain TX5332 transformed either with empty pCIE_spec_ plasmid, or with pCIE_spec_ encoding LsaA. After 16-20 hours at 37 °C without shaking, the presence or absence of bacterial growth was scored by eye.

*B. subtilis* strains were grown in LB medium supplemented with increasing concentrations of antibiotics was inoculated with 5 x 10^5^ CFU/mL (OD_600_ of approximately 0.0005), and after 16-20 hours at 37 °C without shaking the presence or absence of bacterial growth was scored by eye.

### Time-kill kinetics assay

The protocol was based on that of [51] and Svetov [44]. Exponential *L. monocytogenes* cultures in MH-F broth (OD_600_ ≈ 0.3) were diluted to 10^5^ CFU/mL (OD_600_ = 0.001) in 10 mL of MH-F broth either supplemented with appropriate antibiotic at four-fold MIC concentration or without antibiotics (positive growth control), and the resultant cultures were incubated at 37 °C without shaking. 1 mL aliquots were taken at incramental incubation times (0, 2, 4, 6, 8 and 10 h), spun down at 4000 rpm for 5 min at room temperature and cell pellets were gently washed twice with 900 μL of 1x PBS. Cell pellets were resuspended in 100 μL of 1x PBS, ten-fold serial dilutions were prepared in 96-well plates (10^-1^-10^-8^), and 10 μL resultant ten-fold seral dilutions were spotted on BHI agar plates. Colony forming units were scored after 24- to 48-hour incubation at 37 °C.

### Post Antibiotic Effect (PAE) assay

Exponential cultures of *L. monocytogenes* strains in MH-F blood broth media (OD_600_ ≈ 0.3) were diluted to 10^5^ CFU/mL (≈OD_600_ of 0.001) in 5 mL of MH-F media either supplemented with appropriate antibiotic at four-fold MIC concentration or without antibiotics (positive growth control) and incubated at 37 °C for without shaking for 2 h. After the 2 h pre-treatment, antibiotics were removed by 1:100 dilution of 100 μL into 10 mL of fresh prewarmed MH-F blood broth media. At incremental time points (0, 2, 4, 6, 8 and 10 h), 1 mL of the 100x-diluted cell culture was harvested, centrifuged for 5 min at 4000 rpm, 900 μL of the medium was removed, and the pellets were resuspended in the remaining 100 μL. The volume was adjusted to 1 mL with 1x PBS. Control cultures without antibiotics were handled similarly. Cell solutions were then serially diluted ten-fold to 10^-8^, and 10 μL were spotted on BHI agar plates. Plates for individual time points were incubated at room temperature until the last set of plates were spotted (10 h time point), and then incubated at 37 °C. The plates were scored after 24 and 48 h incubation at 37 °C and imaged using ImageQuant LAS 4000 (GE Healthcare). The last time point (24 h) was processed separately analogously to 0-10h time points (see above).

## Acknowledgments

We are grateful to Daniel A. Portnoy for sharing wild-type *L. monocytogenes* 10403S, Jörgen Johansson for sharing wild-type *L. monocytogenes* EGD-e, Barbara E. Murray for sharing *E. faecalis DlsaA* (*lsa*::Kan) strain TX5332 [19] and Birte Vester for sharing the *S. sciuri cfr*-encoding plasmid [50]. This work was supported by the funds from European Regional Development Fund through the Centre of Excellence for Molecular Cell Technology (VH, TT); grant PRG335 from the Estonian Research Council (VH, TT); Swedish Research council (project grants 2017-03783 and 2021-01146, grant 2018-00956 within the RIBOTARGET consortium under the framework of JPIAMR); Ragnar Söderberg foundation (VH). KJYW was supported by a National Science Scholarship (PhD) by the Agency for Science, Technology and Research, Singapore.

## Conflicts of Interest

AGM is an inventor in a provisional patent application submitted by the President and Fellows of Harvard College covering oxepanoprolinamide antibiotics described in this work. AGM has filed the following international patent applications: WO/2019/032936 ‘Lincosamide Antibiotics and Uses Thereof’ and WO/2019/032956 ‘Lincosamide Antibiotics and Uses Thereof’.

## Notes

### Competing Interest Statement

Andrew G. Myers is an inventor in a provisional patent application submitted by the President and Fellows of Harvard College covering oxepanoprolinamide antibiotics described in this work. Andrew G. Myers has filed the following international patent applications: WO/2019/032936 Lincosamide Antibiotics and Uses Thereof and WO/2019/032956 Lincosamide Antibiotics and Uses Thereof.

## References

1 Schwarz S, Shen J, Kadlec K, Wang Y, Brenner Michael G, Fessler AT et al. Lincosamides, Streptogramins, Phenicols, and Pleuromutilins: Mode of Action and Mechanisms of Resistance. Cold Spring Harb Perspect Med 2016; 6.

2 Matzov D, Eyal Z, Benhamou RI, Shalev-Benami M, Halfon Y, Krupkin M et al. Structural insights of lincosamides targeting the ribosome of Staphylococcus aureus. Nucleic Acids Res 2017; 45: 10284–10292.

3 Tu D, Blaha G, Moore PB, Steitz TA. Structures of MLSBK antibiotics bound to mutated large ribosomal subunits provide a structural explanation for resistance. Cell 2005; 121: 257–270.

4 Dunkle JA, Xiong L, Mankin AS, Cate JH. Structures of the Escherichia coli ribosome with antibiotics bound near the peptidyl transferase center explain spectra of drug action. Proc Natl Acad Sci U S A 2010; 107: 17152–17157.

5 Schlunzen F, Zarivach R, Harms J, Bashan A, Tocilj A, Albrecht R et al. Structural basis for the interaction of antibiotics with the peptidyl transferase centre in eubacteria. Nature 2001; 413: 814–821.

6 Spížek J, Řezanka T. Lincosamides: Chemical structure, biosynthesis, mechanism of action, resistance, and applications. Biochem Pharmacol 2017; 133: 20–28.

7 Macleod AJ, Ross HB, Ozere RL, Digout G, Van R. Lincomycin: A New Antibiotic Active against Staphylococci and Other Gram-Positive Cocci: Clinical and Laboratory Studies. Can Med Assoc J 1964; 91: 1056–1060.

8 Birkenmeyer RD, Kagan F. Lincomycin. XI. Synthesis and structure of clindamycin. A potent antibacterial agent. J Med Chem 1970; 13: 616–619.

9 Phillips I. Past and current use of clindamycin and lincomycin. J Antimicrob Chemother 1981; 7 Suppl A: 11–18.

10 Smieja M. Current indications for the use of clindamycin: A critical review. Can J Infect Dis 1998; 9: 22–28.

11 Ahonkhai VI, Cherubin CE, Shulman MA, Jhagroo M, Bancroft U. In vitro activity of U-57930E, a new clindamycin analog, against aerobic gram-positive bacteria. Antimicrob Agents Chemother 1982; 21: 902–905.

12 Birkenmeyer RD, Kroll SJ, Lewis C, Stern KF, Zurenko GE. Synthesis and antimicrobial activity of clindamycin analogues: pirlimycin, a potent antibacterial agent. J Med Chem 1984; 27: 216–223.

13 Hirai Y, Maebashi K, Yamada K, Wakiyama Y, Kumura K, Umemura E et al. Characterization of compound A, a novel lincomycin derivative active against methicillin-resistant Staphylococcus aureus. J Antibiot (Tokyo) 2021; 74: 124–132.

14 Mitcheltree MJ, Pisipati A, Syroegin EA, Silvestre KJ, Klepacki D, Mason JD et al. A synthetic antibiotic class overcoming bacterial multidrug resistance. Nature 2021.

15 Long KS, Poehlsgaard J, Kehrenberg C, Schwarz S, Vester B. The Cfr rRNA methyltransferase confers resistance to Phenicols, Lincosamides, Oxazolidinones, Pleuromutilins, and Streptogramin A antibiotics. Antimicrob Agents Chemother 2006; 50: 2500–2505.

16 Schwarz S, Werckenthin C, Kehrenberg C. Identification of a plasmid-borne chloramphenicol-florfenicol resistance gene in Staphylococcus sciuri. Antimicrob Agents Chemother 2000; 44: 2530–2533.

17 Uchiyama H, Weisblum B. N-Methyl transferase of Streptomyces erythraeus that confers resistance to the macrolide-lincosamide-streptogramin B antibiotics: amino acid sequence and its homology to cognate R-factor enzymes from pathogenic bacilli and cocci. Gene 1985; 38: 103–110.

18 Maravic G. Macrolide resistance based on the Erm-mediated rRNA methylation. Curr Drug Targets Infect Disord 2004; 4: 193–202.

19 Singh KV, Weinstock GM, Murray BE. An Enterococcus faecalis ABC homologue (Lsa) is required for the resistance of this species to clindamycin and quinupristin-dalfopristin. Antimicrob Agents Chemother 2002; 46: 1845–1850.

20 Wilson DN, Hauryliuk V, Atkinson GC, O’Neill AJ. Target protection as a key antibiotic resistance mechanism. Nat Rev Microbiol 2020; 18: 637–648.

21 Murina V, Kasari M, Takada H, Hinnu M, Saha CK, Grimshaw JW et al. ABCF ATPases Involved in Protein Synthesis, Ribosome Assembly and Antibiotic Resistance: Structural and Functional Diversification across the Tree of Life. J Mol Biol 2018.

22 Ero R, Kumar V, Su W, Gao YG. Ribosome protection by ABC-F proteins-Molecular mechanism and potential drug design. Protein Sci 2019; 28: 684–693.

23 Crowe-McAuliffe C, Murina V, Turnbull KJ, Kasari M, Mohamad M, Polte C et al. Structural basis of ABCF-mediated resistance to pleuromutilin, lincosamide, and streptogramin A antibiotics in Gram-positive pathogens. Nat Commun 2021; 12: 3577.

24 Murina V, Kasari M, Hauryliuk V, Atkinson GC. Antibiotic resistance ABCF proteins reset the peptidyl transferase centre of the ribosome to counter translational arrest. Nucleic Acids Res 2018; 46: 3753–3763.

25 Sharkey LK, Edwards TA, O’Neill AJ. ABC-F Proteins Mediate Antibiotic Resistance through Ribosomal Protection. MBio 2016; 7: e01975.

26 Su W, Kumar V, Ding Y, Ero R, Serra A, Lee BST et al. Ribosome protection by antibiotic resistance ATP-binding cassette protein. Proc Natl Acad Sci U S A 2018; 115: 5157–5162.

27 Crowe-McAuliffe C, Graf M, Huter P, Takada H, Abdelshahid M, Novacek J et al. Structural basis for antibiotic resistance mediated by the Bacillus subtilis ABCF ATPase VmlR. Proc Natl Acad Sci U S A 2018; 115: 8978–8983.

28 Radoshevich L, Cossart P. Listeria monocytogenes: towards a complete picture of its physiology and pathogenesis. Nat Rev Microbiol 2018; 16: 32–46.

29 Schlech WF, 3rd, Lavigne PM, Bortolussi RA, Allen AC, Haldane EV, Wort AJ et al. Epidemic listeriosis--evidence for transmission by food. N Engl J Med 1983; 308: 203–206.

30 Southwick FS, Purich DL. Intracellular pathogenesis of listeriosis. N Engl J Med 1996; 334: 770–776.

31 de Noordhout CM, Devleesschauwer B, Angulo FJ, Verbeke G, Haagsma J, Kirk M et al. The global burden of listeriosis: a systematic review and meta-analysis. Lancet Infect Dis 2014; 14: 1073–1082.

32 Mylonakis E, Hohmann EL, Calderwood SB. Central nervous system infection with Listeria monocytogenes. 33 years’ experience at a general hospital and review of 776 episodes from the literature. Medicine (Baltimore) 1998; 77: 313–336.

33 Temple ME, Nahata MC. Treatment of listeriosis. Ann Pharmacother 2000; 34: 656–661.

34 Caruso M, Fraccalvieri R, Pasquali F, Santagada G, Latorre LM, Difato LM et al. Antimicrobial Susceptibility and Multilocus Sequence Typing of Listeria monocytogenes Isolated Over 11 Years from Food, Humans, and the Environment in Italy. Foodborne Pathog Dis 2020; 17: 284–294.

35 Andriyanov PA, Zhurilov PA, Liskova EA, Karpova TI, Sokolova EV, Yushina YK et al. Antimicrobial Resistance of Listeria monocytogenes Strains Isolated from Humans, Animals, and Food Products in Russia in 1950-1980, 2000-2005, and 2018-2021. Antibiotics (Basel) 2021; 10.

36 Rugna G, Carra E, Bergamini F, Franzini G, Faccini S, Gattuso A et al. Distribution, virulence, genotypic characteristics and antibiotic resistance of Listeria monocytogenes isolated over one-year monitoring from two pig slaughterhouses and processing plants and their fresh hams. Int J Food Microbiol 2021; 336: 108912.

37 Tirziu E, Herman V, Nichita I, Morar A, Imre M, Cucerzan A et al. Diversity and antibiotic resistance profiles of Listeria monocytogenes serogroups in different food products from Transylvania Region, Central Romania. J Food Prot 2021.

38 Dar D, Shamir M, Mellin JR, Koutero M, Stern-Ginossar N, Cossart P et al. Term-seq reveals abundant ribo-regulation of antibiotics resistance in bacteria. Science 2016; 352: aad9822.

39 Ohki R, Tateno K, Takizawa T, Aiso T, Murata M. Transcriptional termination control of a novel ABC transporter gene involved in antibiotic resistance in Bacillus subtilis. J Bacteriol 2005; 187: 5946–5954.

40 Glaser P, Frangeul L, Buchrieser C, Rusniok C, Amend A, Baquero F et al. Comparative genomics of Listeria species. Science 2001; 294: 849–852.

41 Edman DC, Pollock MB, Hall ER. Listeria monocytogenes L forms. I. Induction maintenance, and biological characteristics. J Bacteriol 1968; 96: 352–357.

42 Bécavin C, Bouchier C, Lechat P, Archambaud C, Creno S, Gouin E et al. Comparison of widely used Listeria monocytogenes strains EGD, 10403S, and EGD-e highlights genomic variations underlying differences in pathogenicity. mBio 2014; 5: e00969–00914.

43 Britton RA, Eichenberger P, Gonzalez-Pastor JE, Fawcett P, Monson R, Losick R et al. Genome-wide analysis of the stationary-phase sigma factor (sigma-H) regulon of Bacillus subtilis. J Bacteriol 2002; 184: 4881–4890.

44 Svetlov MS, Vazquez-Laslop N, Mankin AS. Kinetics of drug-ribosome interactions defines the cidality of macrolide antibiotics. Proc Natl Acad Sci U S A 2017; 114: 13673–13678.

45 Walkup GK, You Z, Ross PL, Allen EK, Daryaee F, Hale MR et al. Translating slow-binding inhibition kinetics into cellular and in vivo effects. Nat Chem Biol 2015; 11: 416–423.

46 Bundtzen RW, Gerber AU, Cohn DL, Craig WA. Postantibiotic suppression of bacterial growth. Rev Infect Dis 1981; 3: 28–37.

47 Mohamad M, Nicholson D, Saha CK, Hauryliuk V, Edwards TA, Atkinson GC et al. Sal-type ABC-F proteins: intrinsic and common mediators of pleuromutilin resistance by target protection in staphylococci. Nucleic Acids Res 2022.

48 Sharkey LKR, O’Neill AJ. Antibiotic Resistance ABC-F Proteins: Bringing Target Protection into the Limelight. ACS Infect Dis 2018; 4: 239–246.

49 Mason JD, Terwilliger DW, Pote AR, Myers AG. Practical Gram-Scale Synthesis of Iboxamycin, a Potent Antibiotic Candidate. J Am Chem Soc 2021; 143: 11019–11025.

50 Ntokou E, Hansen LH, Kongsted J, Vester B. Biochemical and Computational Analysis of the Substrate Specificities of Cfr and RlmN Methyltransferases. PLoS One 2015; 10: e0145655.

51 Barry AL, Craig WA, Nadler H, Reller BL, Sanders CC, Swenson JM. Methods for Determining Bactericidal Activity of Antimicrobial Agents; Approved Guideline. CLSI document M26-A Waybe, PA: Clinical and Laboratory Standarts Institute 1999.

